# A novel Fiji/ImageJ plugin for the rapid analysis of blebbing cells

**DOI:** 10.1101/2021.06.04.447112

**Authors:** Karl W. Vosatka, Sandrine B. Lavenus, Jeremy S. Logue

## Abstract

When confined, cells have recently been shown to undergo a phenotypic switch to what has been termed, fast amoeboid (leader bleb-based) migration. However, as this is a nascent area of research, few tools are available for the rapid analysis of cell behavior. Here, we demonstrate that a novel Fiji/ImageJ-based plugin, Analyze_Blebs, can be used to quickly obtain cell migration parameters and morphometrics from time lapse images. As validation, we show that Analyze_Blebs can detect significant differences in cell migration and morphometrics, such as the largest bleb size, upon introducing different live markers of F-actin, including F-tractin and LifeAct tagged with green and red fluorescent proteins. We also demonstrate, using flow cytometry, that live markers increase total levels of F-actin. Furthermore, that F-tractin increases cell stiffness, which was found to correlate with a decrease in migration, thus reaffirming the importance of cell mechanics as a determinant of Leader Bleb-Based Migration (LBBM).

**Highlight summary:** A new plugin, Analyze_Blebs, enables the rapid analysis of cell migration and morphometrics of fast amoeboid cells. Morphometrics combined with cell stiffness are found to be predictive of confined, fast amoeboid (leader bleb-based), migration.

## Introduction

Cell migration mediates diverse physiological processes, including embryonic development, immune surveillance, and wound healing. Accordingly, the de-regulation of cell migration is a hallmark of disease. A large portion of cell migration research has been conducted on glass substrates, which favors (integrin-dependent) mesenchymal motility. Under conditions that more closely resemble the tissue environment, cells have been shown to rapidly switch between a growing list of migration modes [1, 2], optimizing their migratory potential based on their immediate (tissue) environment. Under conditions of high (mechanical) confinement, a diverse range of cell types have recently been shown to undergo a phenotypic transition to fast amoeboid or Leader Bleb-Based Migration (LBBM) [3-6].

A hallmark of amoeboid migration is the formation of Plasma Membrane (PM) blebs. Blebs form in response to a local separation between the PM and underlying cortical actin cytoskeleton [7]. Accordingly, cells with elevated intracellular pressure often display numerous blebs [8]. In tissue culture, blebs are normally dynamic – forming and retracting in less than a minute [7]. However, when cells are confined down to 3 μm (which has been found to be optimal) they may form a very large and stable bleb [4]. Within these stable blebs, cortical actin flows from the bleb tip to the neck, which is enriched with myosin and separates the bleb from the cell body [3-6]. Importantly, with non-specific friction, it is this cortical actin flow that provides the motive force for cell movement [6]. Accordingly, as cells move in the direction of this bleb, it has been termed a ‘leader bleb’ [3]. The hallmarks of LBBM have also been observed *in vivo*, including within developing Zebrafish embryos and tumors by intravital imaging [5, 9]. Thus, LBBM is an important new mode of migration that cells may adopt under confinement.

The study of LBBM presents a quantitative challenge. Bleb dynamics are fast compared to the speed of migration. Blebs can form and retract in less than a minute, whereas cells displaying LBBM migrate at a speed of about 30 μm per hour [3-6]. Other plugins for analyzing blebs rely on a characteristic change in curvature [16-17]. Due to their unique shape, leader blebs are not likely to be accurately identified by this method. Moreover, these plugins require very high temporal resolution for tracking individual blebs [16-17]. In order to balance the risk of photobleaching and cell damage against the speed of both migration and bleb formation/retraction, it is necessary to target a “Goldilocks” time resolution for live imaging. Our lab has had success imaging cells for several hours with images every few minutes, which allows for accurate capturing of motility and, in some cases, time for cells to switch to LBBM. This paradigm, as well as any other that tries to capture a significant picture of cell motility, makes it difficult to capture bleb dynamics; blebs appear and disappear between frames, making it very difficult for a computer to accurately track them.

Yet another challenge in this nascent research area is in deciding which parameters to employ in understanding cell motility. The molecular mechanism(s) that control LBBM are not fully understood. In melanoma cells, it is clear that the actin bundling and capping protein, Eps8, reduced Src tyrosine kinase activity, and a compliant network of Vimentin are essential to LBBM [3, 10, 11]. Thus, in studying the behavior of confined cells, it may not be clear what characteristics of blebbing are most important. There is a wide range of properties of blebbing cells that may be of interest, including the number of blebs, bleb area, leader bleb area, and cell body area.

Manual analysis of LBBM requires precise tracing of the outline of the whole cell and sub-cellular elements like blebs. Time-lapses are by necessity long and require multiple sessions of tracing to capture sub-cellular components separately from whole cell measurements. The process is further complicated by the frequent fluctuation in cell shape and the dynamic occurrence of blebs. Consequently, user input into the analysis process is often necessary because blebs are very difficult to characterize without user input. Their highly dynamic nature makes them difficult for computers to track, and they cannot be easily stained as separate from the cell body. Still, a fully manual analysis process, in addition to being quite slow, is subject to increased user bias and lower reproducibility.

Here, we present a custom plugin for the imaging analysis software, Fiji/ImageJ (NIH), for use in studies of LBBM. The plugin is a manual-auto hybrid that employs image thresholding techniques and automatic quantification to drastically increase analysis speed. The plugin is supervised by the user and takes advantage of the ease of identifying blebs by eye. It offers a solution that can be used for even the noisiest imaging time courses, as it allows for manual tracing of the cell outline. This allows for consistent and highly reproducible analysis of time course images. As validation, we used this plugin to compare the effects of different live cell markers of F-actin on cell behavior. We report that a subset of these markers can significantly impede confined (leader bleb-based) migration. Moreover, a subset of these markers led to a decrease in cell compressibility. Thus, we provide a valuable new resource for the rapid analysis of LBBM.

## Results

### Mechanically confined melanoma cells adopt three distinct phenotypes

For the mechanical confinement of cells, we previously developed an assay that places cells underneath a slab of PDMS held at a defined height above cover glass by micron-sized beads (Fig. 1A) [12]. Previously, it was shown that the phenotypic transition to fast amoeboid (leader bleb-based) migration occurs most efficiently when cells are confined down to 3 μm [4]. Moreover, it has been shown that fast amoeboid cells transmit motive forces through non-specific friction, which can be modulated through surface coatings [6]. In our confinement assay, all surfaces are coated with 1% BSA to increase non-specific friction. Although Liu *et al*. demonstrated that several cell types can undergo this phenotypic transition, melanoma cells represent the best studied model of LBBM. Therefore, we have used melanoma A375-M2 cells for validating the Fiji/ImageJ-based plugin described here.

**Figure1.**
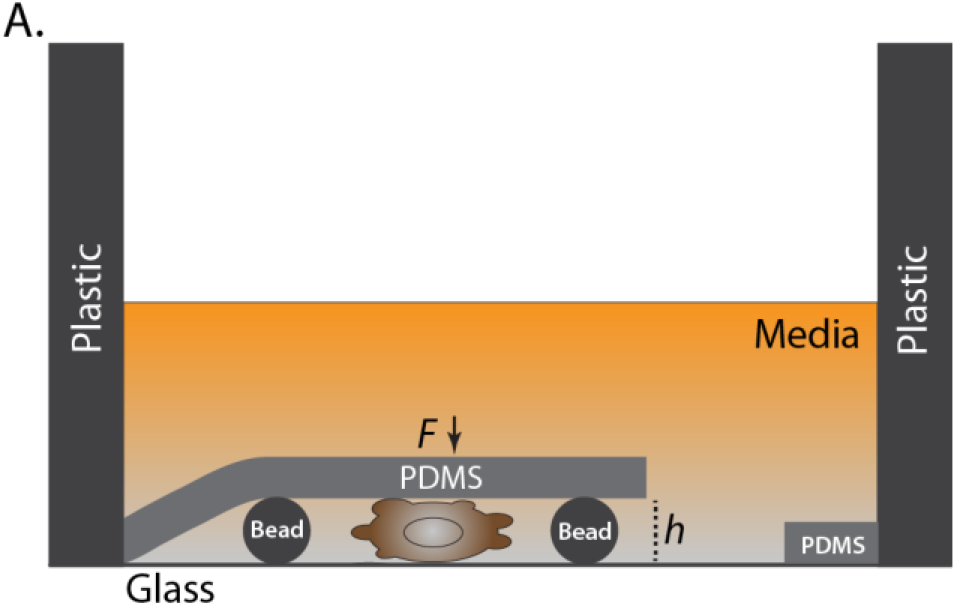
Schematic of the 2D confinement assay. **A**. The confinement height (*h*) is maintained by micron-sized beads. Downward force (*F*) onto the cells and beads comes from the attachment between the elastic PDMS and plastic wall of a glass-bottom well or tissue culture dish.

When A375-M2 cells are confined down to 3 μm, we observe three distinct phenotypes. First, we observe what we refer to as the leader mobile (LM) population, which undergoes rapid (directional) LBBM (Fig. 2A; *top*). Second, we observe what we refer to as the leader non-mobile (LNM) population, which are non-mobile cells with one large and stable bleb (Fig. 2A; *middle*). Our previous work has demonstrated that LNM cells are in essence stuck, as stiffening of the Vimentin network was shown to increase the proportion of LNM cells [11]. Finally, there is a population of cells we refer to as no leader (NL) cells, which are non-mobile cells that form many dynamic circumferential blebs (Fig. 2A; *bottom*). A given perturbation may lead to changes in the relative number of each phenotype; therefore, deciphering the underlying mechanisms for each change requires detailed morphometric analyses.

**Figure 2.**
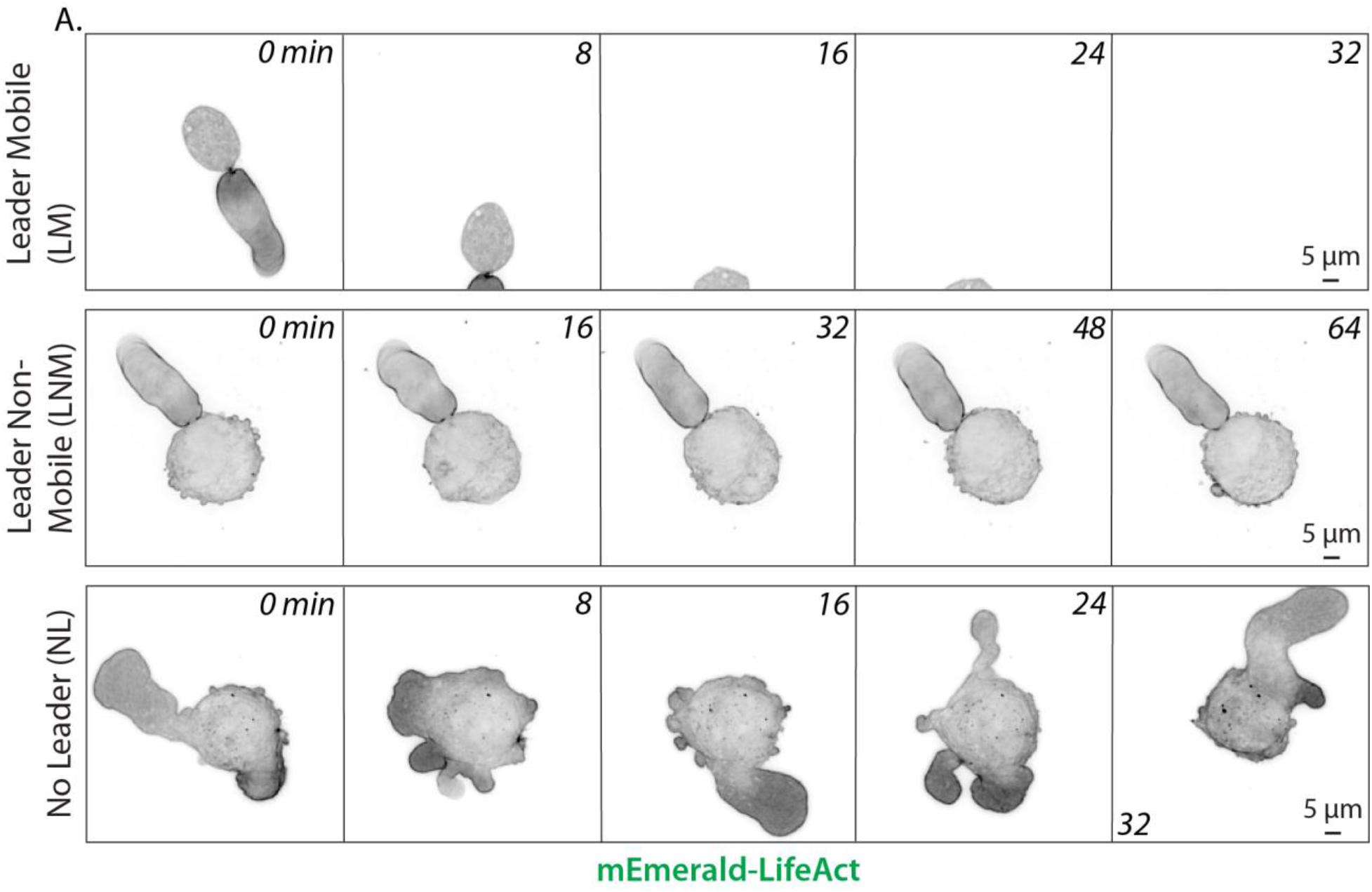
Mechanically confined melanoma cells adopt three distinct phenotypes. **A**. When confined down to 3 μm, melanoma A375-M2 cells adopt one of three phenotypes, including leader mobile (LM; form a leader bleb and migrate), leader non-mobile (LNM; form a leader bleb but do not migrate), and no leader (NL; do not form a leader bleb and do not migrate) phenotypes. All cells were transfected with the F-actin marker, LifeAct-mEmerald, before confinement. All data are representative of at least three independent experiments.

### Overview of the Analyze_Blebs plugin

The Fiji/ImageJ-based plugin, Analyze_Blebs, begins by prompting users to indicate a name and save path for plugin files. Representative images for each stage of the plugin are saved in the chosen file path, along with a Comma Separated Values (CSV) file compatible with Excel (Microsoft, Redmond, WA) containing data corresponding to each plugin-generated image. Users are also given the option to save data from each plugin-generated image from one cell together in one sheet of an Excel file. Then, users isolate a single stack of images and standardize the image input for proper data analysis. First, if multiple color channels are present, the preferred channel for the analysis process is selected. Then, the image is isolated to a single Z stack, either by selecting one Z slice from those present or by choosing a method for collapsing stacks such as a maximum intensity projection (MIP). Finally, if the image is scaled to pixels, the user is prompted to convert the image to the micron scale. Image analysis measurement options, if they differ from the plugin’s default options, can also be chosen at this stage. The plugin will automatically load the last set of measurement options used by the plugin.

The image then is further edited to ensure the cell of interest can be isolated. In the first stage of image editing, users are prompted to edit their raw data file to erase any objects on the scale of the cell that appear in the background that might interfere with thresholding methods; this includes removing all cells present in the image besides the one to be analyzed. The resulting edited file is saved by the default name “Tracer.tif”. This edited raw image is then binarized using image thresholding approaches (Fig. 3A). All image thresholding options are run on the first frame of the time stack and presented to the user to select from. The user can choose a method, preview its results through the time stack, and either move forward with the chosen threshold or return to the preview stage and pick a different one. This initial binary image is saved as “Threshold.tif” with the thresholding method appended to the file name for later reference. The user then has the option to further edit the threshold if desired, by first filling any holes within the object of interest, removing any background images with size exclusion and removing any noise objects attached to the image (Fig. 3A). The resulting cleaned binary image is saved as a “Whole_cell.tif” image. Finally, the plugin automatically creates a series of binary image stacks with isolated subcellular structures (Fig. 3B). The user is prompted to circle the cell body on each frame, after which it is removed. This process yields an image with just blebs (“All_blebs.tif”), which is subtracted from the Whole_cell image to isolate the cell body (“Cell_body.tif”). The plugin also calculates an image series from the All_blebs images that automatically isolates the largest bleb by area on each frame of the time series (“Largest_blebs.tif”). Data files are automatically created to accompany each image and can be included together in one worksheet in an optional Excel summary document. Other options include the ability to restart the process partway by running the plugin with the output images and the ability to automatically analyze multiple cells on the same image series. A more detailed protocol with instructions and tips for use of the plugin is included with Supporting Information.

**Figure 3.**
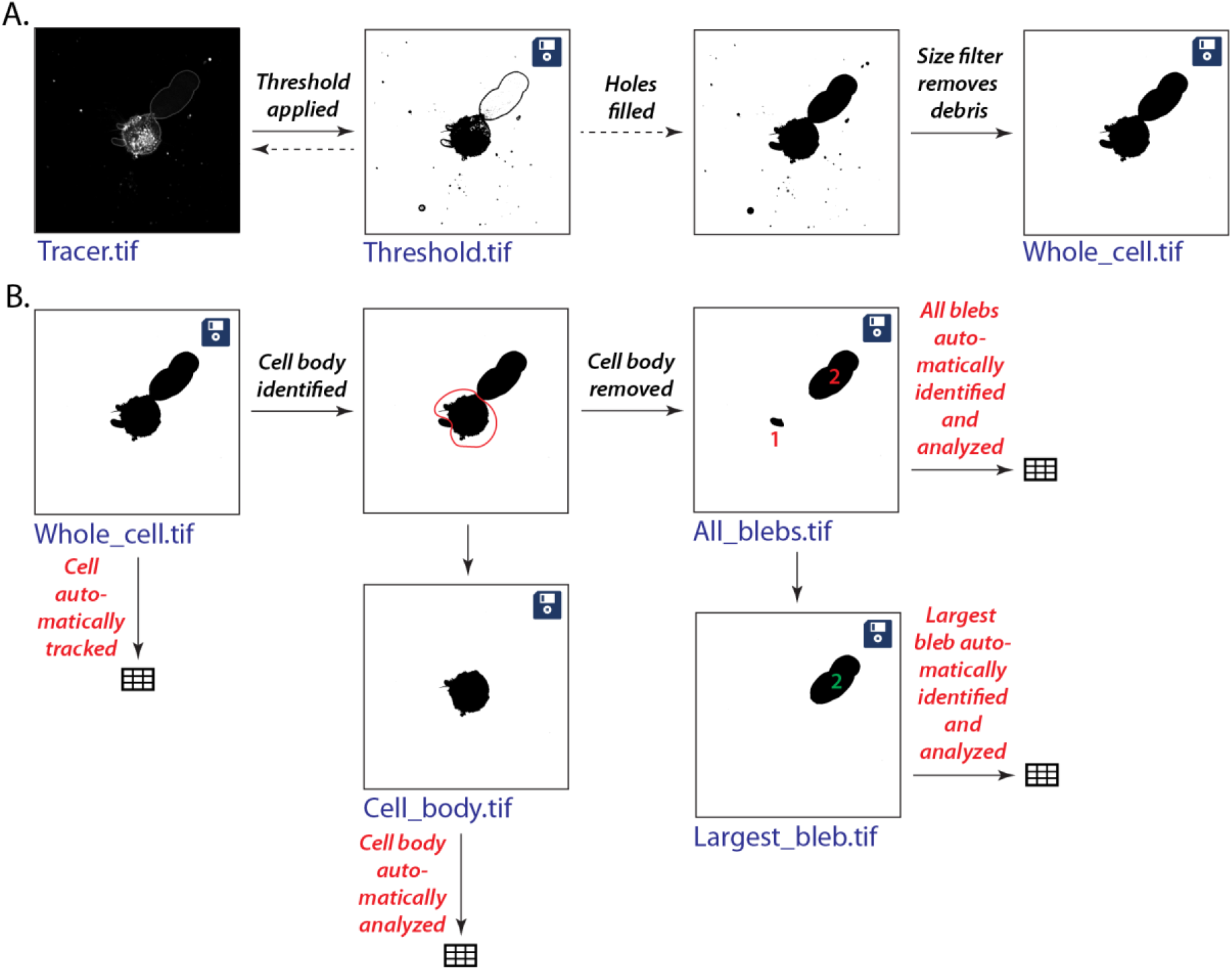
Overview of the Analyze_Blebs plugin. **A**. Image binarization approach is previewed and selected. The threshold choice can be edited further to fill holes and remove any attached noise. **B**. Analysis workflow creates a series of binary image stacks with isolated subcellular structures. Whole cell is first isolated with size exclusion, then the cell body is removed by user input, and remaining images are created by the plugin. Disk icon on image indicates that the image series is saved with the name listed in blue. Spreadsheet icon indicates a CSV file was saved for that image analysis under image name.

**Figure 4.**
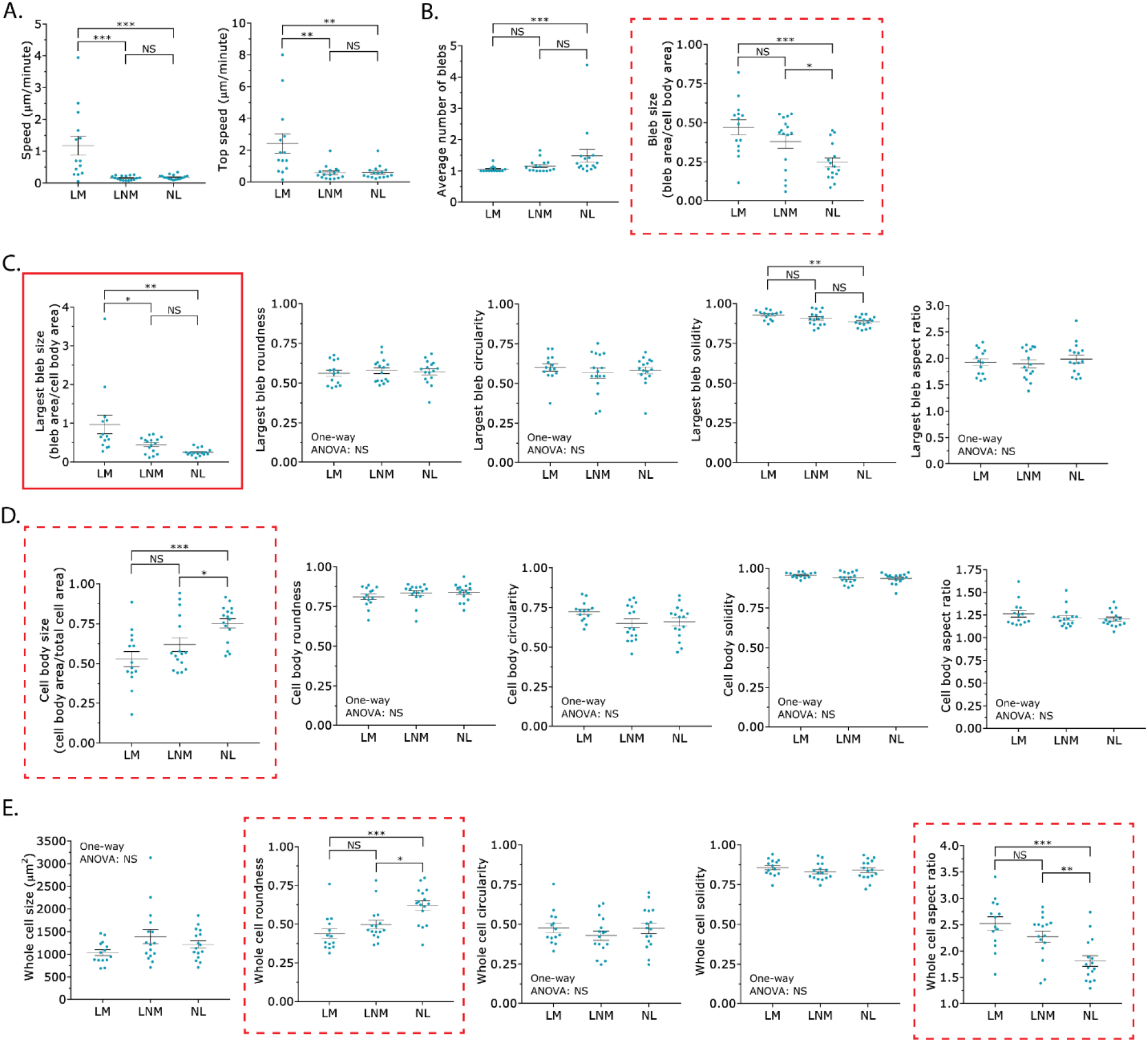
Shape descriptors may be used to characterize each phenotype. **A**. Instantaneous speeds (*left*) and top instantaneous speeds (*right*) for each phenotype. **B**. Average number of blebs (*left*) and bleb size (*right*) for each phenotype. **C**. Shape descriptors for the largest bleb across each phenotype. Solid red outline highlights the average largest bleb size as a shape descriptor that can differentiate between leader mobile (LM) from leader non-mobile (LNM) cells. **D**. Shape descriptors for the cell body across each phenotype. Dashed red outline highlights the average cell body size as a shape descriptor that can differentiate between leader non-mobile (LNM) from no leader (NL) cells. **E**. Shape descriptors for the whole cell across each phenotype. Dashed red outline highlights average whole cell roundness and average whole cell aspect ratio as shape descriptors that can differentiate between leader non-mobile (LNM) and no leader (NL) cells. Error is SEM. Statistical significance was determined using a One-way ANOVA and a Tukey’s post hoc test. All data are representative of at least three independent experiments. * - p ≤ 0.05, ** - p ≤ 0.01, *** - p ≤ 0.001, and **** - p ≤ 0.0001

### Shape descriptors may be used to characterize each phenotype

The identification of specific shape descriptors that correlate with LBBM is important because it may i) provide novel insight into mechanisms of LBBM and ii) facilitate the use of High Content Screening (HCS) approaches. Therefore, using Analyze_Blebs, we determined if specific shape descriptors, including 1) the average number of blebs, 2) bleb size, 3) largest bleb size, 4) largest bleb roundness, 5) largest bleb circularity, 6) largest bleb solidity, 7) largest bleb aspect ratio, 8) cell body size, 9) cell body roundness, 10) cell body circularity, 11) cell body solidity, 12) cell body aspect ratio, 13) whole cell size, 14) whole cell roundness, 15) whole cell circularity, 16) whole cell solidity, and 17) whole cell aspect ratio correlated with either the leader mobile (LM), leader non-mobile (LNM), or no leader (NL) phenotypes. Besides using speed, a challenge in using shape descriptors is finding one that can discriminate between the leader mobile (LM) and leader non-mobile (LNM) populations, as these cells are phenotypically very similar (Fig. 2A). In addition to differentiating between the leader mobile (LM) and no leader (NL) populations, 4 shape descriptors, including the average 1) bleb size, 2) cell body size, 3) whole cell roundness, and 4) whole cell aspect ratio could differentiate between the leader non-mobile (LNM) and no leader (NL) populations (Fig. 2B-E; *red dashed outlines*). The average largest bleb solidity could only differentiate between the leader mobile (LM) and no leader (NL) populations (Fig. 2C). Only one shape descriptor, the average largest bleb size, could differentiate between the leader mobile (LM) and leader non-mobile (LNM) populations (Fig. 2C; *red solid outline*). Additionally, the average largest bleb size could differentiate between the leader mobile (LM) and no leader (NL) populations (Fig. 2C; *red solid outline*). Thus, these data would support the use of the largest bleb size as a meaningful shape descriptor.

### Analyze_Blebs identifies the limitations of F-actin markers in studying the behavior of fast amoeboid cells

Fast amoeboid (leader bleb-based) migration relies on the coordinated assembly and disassembly of the F-actin cytoskeleton. Therefore, we used Analyze_Blebs to systematically compare the ability of the live markers, F-tractin and LifeAct, to label F-actin without perturbing cell behavior. Using green and red fluorescent protein fusions, both F-tractin and LifeAct were found to mark the cortical actin cytoskeleton in leader blebs and cell body (Fig. 5A). We then used time lapse imaging to determine if F-tractin and/or LifeAct affected LBBM. As a first approach, we determined the percent of leader mobile (LM), leader non-mobile (LNM), and no leader (NL) cells upon staining with a membrane dye only, introduction of EGFP alone, EGFP-F-tractin, mEmerald alone, and mEmerald-LifeAct. This analysis revealed that the number of leader mobile (LM) cells was reduced upon transient transfection of EGFP-F-tractin, whereas the other conditions only moderately effected this number (Fig. 6A). Strikingly, the introduction of EGFP alone or mEmerald alone significantly increased (>50%) speeds for all cells (Fig. 6B). Compared to membrane dye alone, all conditions showed an increase in the speed of the leader mobile (LM) population; however, only mEmerald alone reached statistical significance with an over 3-fold increase in speed (Fig. 6B’). Subsequently, we analyzed cells with red fluorescent protein tags, including FusionRed alone, FusionRed-F-tractin, mScarlet alone, and mScarlet-LifeAct. Again, upon calculating the percent of each population, we found that transfection of Fusion-F-tractin reduced the number of leader mobile (LM) cells (Fig. 6C). Surprisingly, the introduction of FusionRed alone had a similar effect, whereas the other conditions had similar numbers of leader mobile (LM), leader non-mobile (LNM), and no leader (NL) cells (Fig. 6C). In line with a decrease in the number of leader mobile (LM) cells, the introduction of FusionRed-F-tractin significantly decreased speeds (>50%) for all cells (Fig. 6D). In contrast, we found that transfection of mScarlet alone and mScarlet-LifeAct increased speeds, however, only mScarlet-LifeAct reached statistical significance (Fig. 6D). Compared to membrane dye only, the introduction of FusionRed alone, mScarlet alone, and mScarlet-LifeAct increased speeds for the leader mobile (LM) population, however, none of these increases reached statistical significance (Fig. 6D’). Due to the very low number of leader mobile (LM) cells for the FusionRed-F-tractin condition, we could not calculate speeds (Fig. 6D’). Thus, these results reveal that both F-tractin and LifeAct, in addition to the fluorescent protein tag, can significantly affect the behavior of mechanically confined cells.

**Figure 5.**
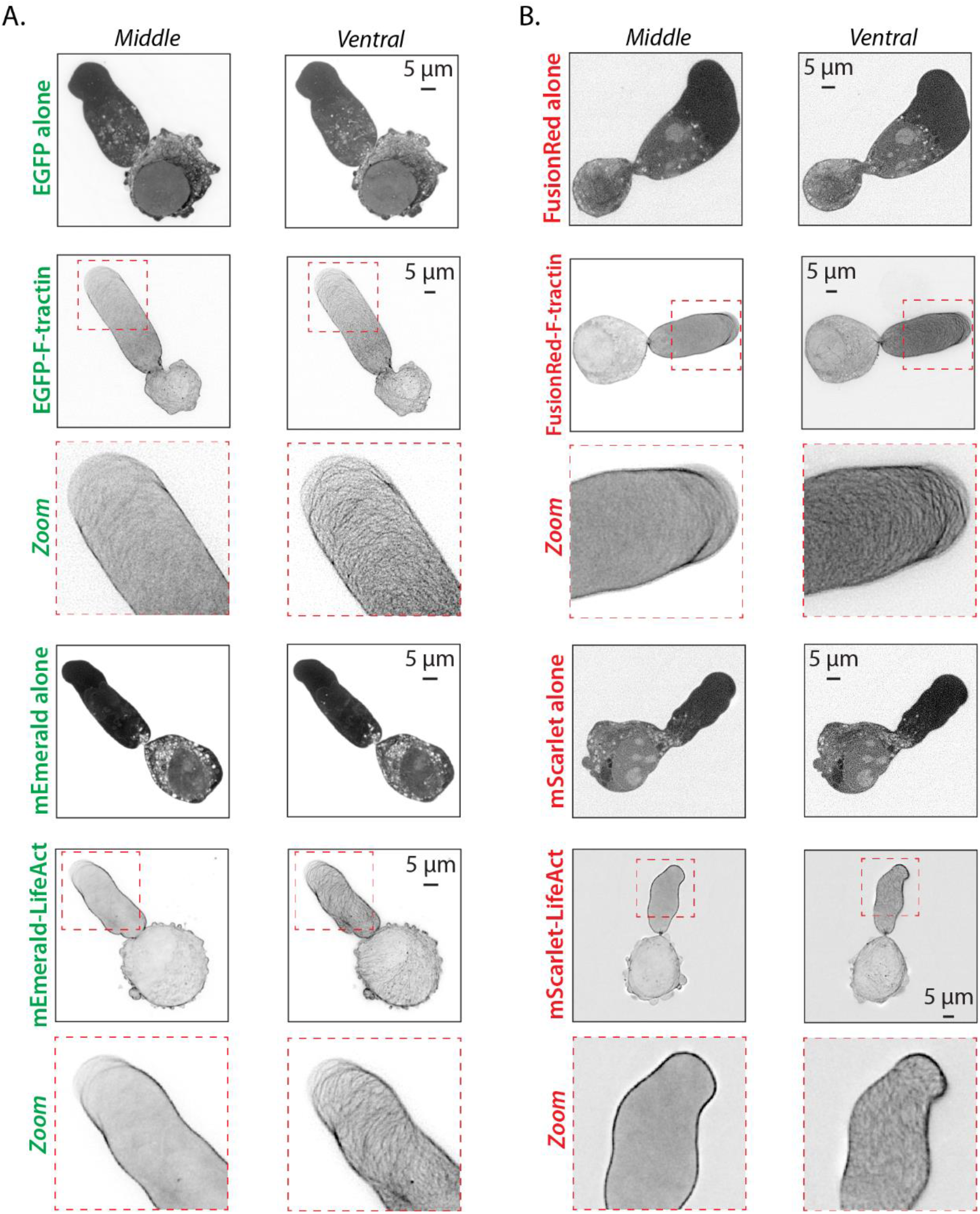
F-tractin and LifeAct mark the cortical actin cytoskeleton in leader blebs. **A**. Middle and ventral Z-sections of melanoma A375-M2 cells transiently transfected with EGFP alone, EGFP-F-tractin, mEmerald alone, and mEmerald-LifeAct. Zooms highlight the intricate cortical actin network in leader blebs. **B**. Middle and ventral Z-sections of melanoma A375-M2 cells transiently transfected with FusionRed alone, FusionRed-F-tractin, mScarlet alone, and mScarlet-LifeAct. Zooms highlight the intricate cortical actin network in leader blebs. All data are representative of at least three independent experiments.

**Figure 6.**
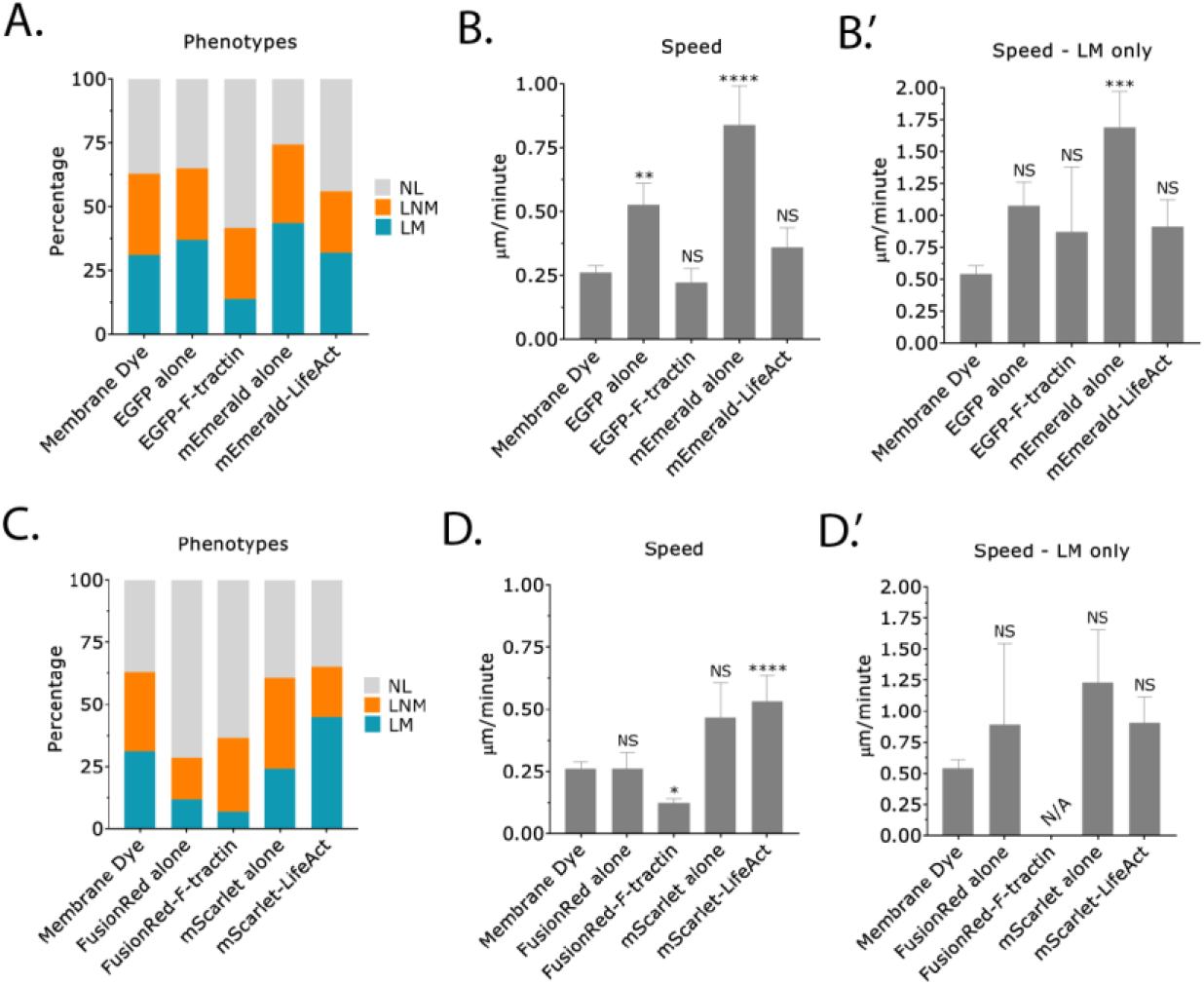
LBBM is affected by fluorescent protein markers. **A**. Percent of no leader (NL), leader non-mobile (LNM), and leader mobile (LM) for melanoma A375-M2 cells stained with membrane dye (n=124 cells), transiently transfected with EGFP alone (n=92 cells), EGFP-F-tractin (n=43 cells), mEmerald alone (n=37 cells), or mEmerald-LifeAct (n=47 cells). **B**. Instantaneous speeds for all cells transfected as in (*A*). **B’**. Instantaneous speeds for leader mobile (LM) cells transfected as in (*A*). **C**. Percent of no leader (NL), leader non-mobile (LNM), and leader mobile (LM) for melanoma A375-M2 cells stained with membrane dye (n=124 cells), transiently transfected with FusionRed alone (n=45 cells), FusionRed-F-tractin (n=29 cells), mScarlet alone (n=33 cells), or mScarlet-LifeAct (n=47 cells). **D**. Instantaneous speeds for all cells transfected as in (*C*). **D’**. Instantaneous speeds for leader mobile (LM) cells transfected as in (*C*). Error is SEM. Statistical significance was determined using a One-way ANOVA and a Kruskal-Wallis post hoc test. All data are representative of at least three independent experiments. * - p ≤ 0.05, ** - p ≤ 0.01, *** - p ≤ 0.001, and **** - p ≤ 0.0001

Live markers of F-actin have previously been reported to perturb a number of cellular processes, including embryonic development [13]. Therefore, we utilized Analyze_Blebs to determine in a high-throughput fashion how F-tractin, LifeAct, and fluorescent proteins tags might affect cell morphometrics. Using the previously described shape descriptors, we found that the transfection of several proteins, including EGFP-F-tractin and FusionRed-F-tractin (which inhibit migration; Fig 6), led to statistically significant increases in the largest bleb area (Fig. 7C). In contrast, we could not detect any changes in the average number of blebs, bleb size, cell body size, whole cell aspect ratio, or whole cell roundness (Fig. 7A-B & D-F). Because the largest bleb size usually positively correlates with cell migration, we were surprised by this result; therefore, we set out to determine if these markers perturb normal F-actin dynamics. To accomplish this, we used a flow cytometry based assay in which the total level of F-actin could be determined in many transfected cells, as measured by the incorporation of fluorescently conjugated phalloidin. The specificity of the approach was confirmed by treating cells with 5 μM Latrunculin-A (Lat-A; 10 min), which reduced the total level of F-actin by over 75% (relative to untreated cells; Fig. 8A). Interestingly, transfection of EGFP-F-tractin and mEmerald-LifeAct led to a roughly 25% increase in F-actin, which is in line with previous studies showing that F-actin may be stabilized by side binding proteins (Fig. 8A) [14]. Similarly, transfection of FusionRed-F-tractin and mScarlet-LifeAct led to significant increases in F-actin (Fig. 8B). Strikingly, the transfection of FusionRed alone also increased the total level of F-actin (Fig. 8B). Although the observed increases are in line with the notion that side binding proteins stabilize F-actin, these results do not fully correlate with the observed decreases in the rate of LBBM. Thus, in order to better understand this phenomenon, we turned to measuring cell stiffnesses. This was done using a previously described assay in which cells are compressed between two polyacrylamide (PA) gels of known stiffness (1 kPa) [4]. A ratio of the cell height to the width (*h*/*d*) is used to define the cell stiffness (Fig. 8C). Using this approach, we found that the transfection of EGFP-F-tractin led to a statistically significant increase in cell stiffness, whereas the transfection of EGFP alone, mEmerald alone, and mEmerald-LifeAct had no effect (relative to membrane dye; Fig. 8D). Similarly, the transfection of FusionRed alone and FusionRed-F-tractin led to a statistically significant increase in cell stiffness, whereas mScarlet-LifeAct had no effect (Fig. 8E). In contrast, the transfection of mScarlet alone led to a statistically significant decrease in cell stiffness (Fig. 8E). Thus, an increase in stiffness perfectly correlates with a decrease in the rate of LBBM for melanoma cells transfected with fluorescent protein markers.

**Figure 7.**
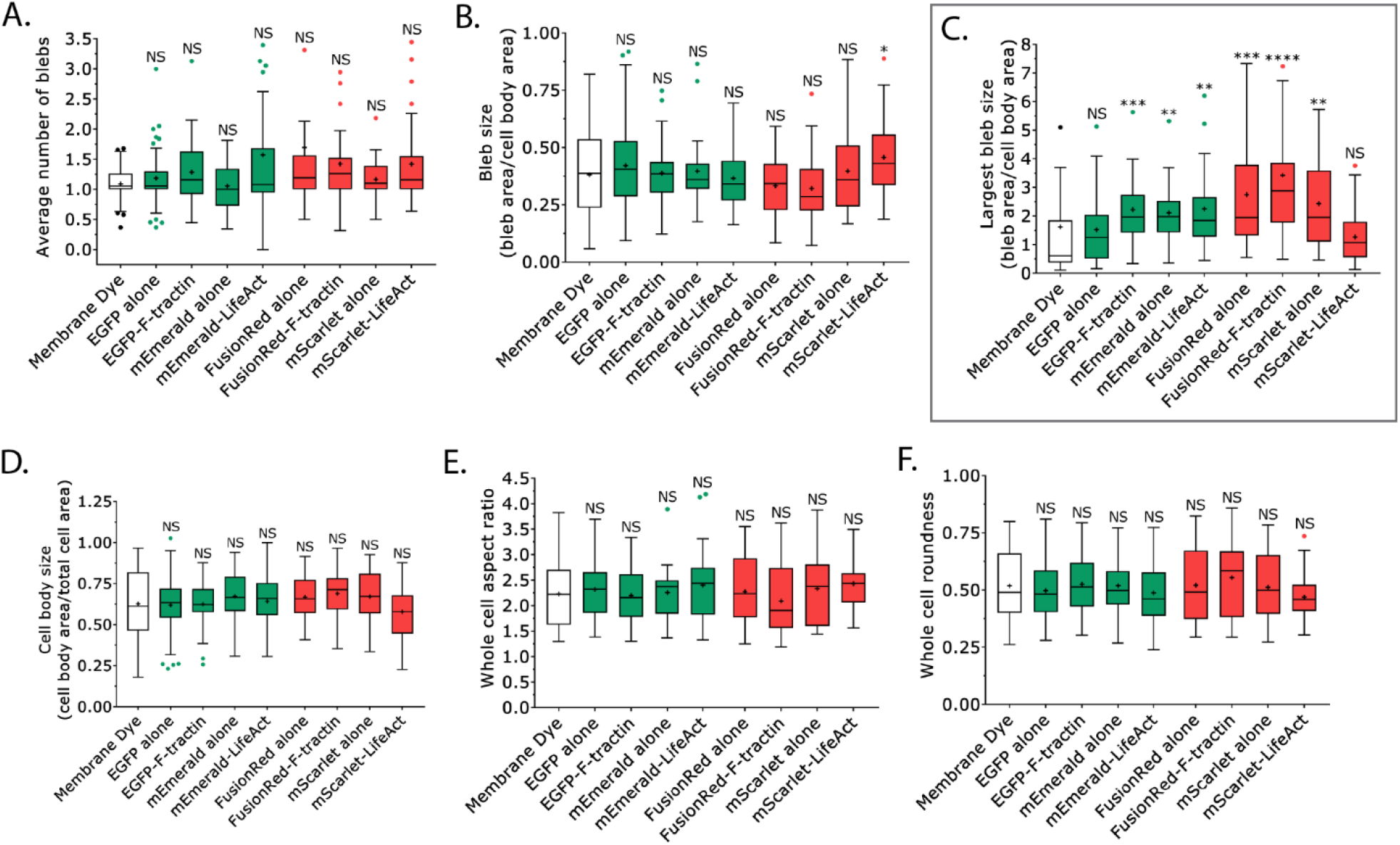
Cell morphometrics are affected by fluorescent protein markers. **A-F**. Using Analyze_Blebs, the average number of blebs (*A*), bleb size (all blebs; *B*), largest bleb size (*C*), cell body size (*D*), whole cell aspect ratio (*E*), and whole cell roundness (*F*) was determined for melanoma A375-M2 cells stained with membrane dye (n=35 cells), transiently transfected with EGFP alone (n=82 cells), EGFP-F-tractin (n=43 cells), mEmerald alone (n=27 cells), mEmerald-LifeAct (n=43 cells), FusionRed alone (n=40 cells), FusionRed-F-tractin (n=28 cells), mScarlet alone (n=30 cells), or mScarlet-LifeAct (n=51 cells) (“+” and line denote the mean and median, respectively). Statistical significance was determined using a One-way ANOVA and a Tukey’s post hoc test. All data are representative of at least three independent experiments. * - p ≤ 0.05, ** - p ≤ 0.01, *** - p ≤ 0.001, and **** - p ≤ 0.0001

**Figure 8.**
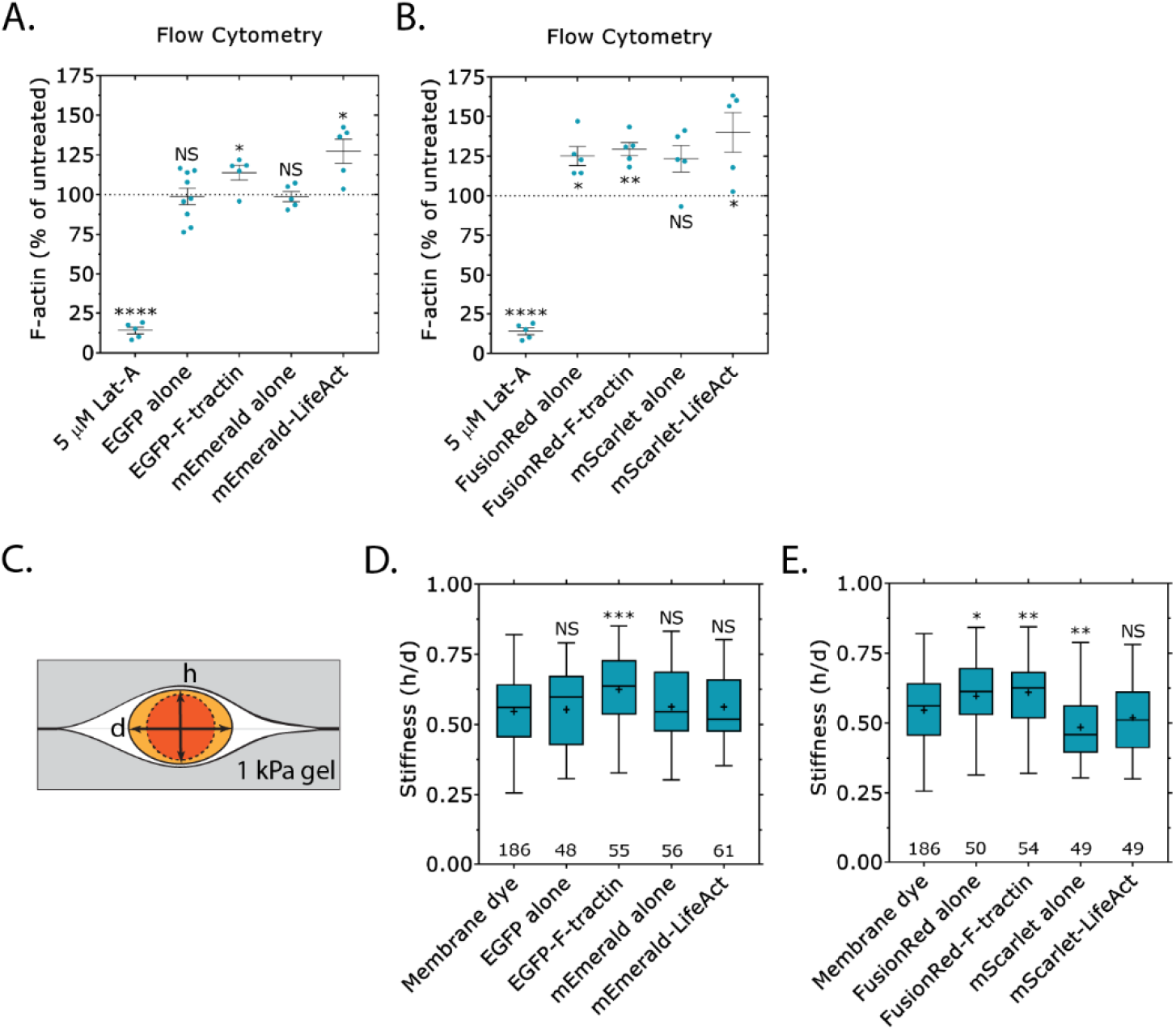
F-actin levels and cell stiffness is affected by fluorescent protein markers. **A-B**. Using a flow cytometry based assay, the total levels of F-actin were determined in freshly trypsinized (spherical) melanoma A375-M2 cells after transient transfection with EGFP alone, EGFP-F-tractin, mEmerald alone, mEmerald-LifeAct, mEmerald alone, mEmerald-LifeAct (*A*) or FusionRed alone, FusionRed-F-tractin, mScarlet alone, and mScarlet-LifeAct (*B*). Treatment with 5 μM Latrunculin-A (Lat-A; 10 min) confirms the specificity of the approach. All data are normalized to untreated controls. **C**. Schematic representation of the cell stiffness assay in which cells are sandwiched between polyacrylamide (PA) gels of know stiffness (1 kPa). A ratio of the cell height (*h*) to the diameter (*d*) is used to define the cell stiffness. **D-E**. Using the cell stiffness assay shown in (*C*), the stiffness of melanoma A375-M2 cells transiently transfected with EGFP alone, EGFP-F-tractin, mEmerald alone, mEmerald-LifeAct (*D*) or FusionRed alone, FusionRed-F-tractin, mScarlet alone, and mScarlet-LifeAct (*E*) (“+” and line denote the mean and median, respectively). Statistical significance was determined using a One-way ANOVA and a Tukey’s post hoc test. All data are representative of at least three independent experiments. * - p ≤ 0.05, ** - p ≤ 0.01, *** - p ≤ 0.001, and **** - p ≤ 0.0001

Diverse cell types form plasma membrane and nuclear blebs under a variety of conditions, including under confinement, hypotonic shock, and apoptosis. During apoptosis, blebbing is observed before cells are orderly disassembled into apoptotic bodies [15]. Therefore, we wondered if Analyze_Blebs could be used to distinguish differences in cell morphometrics between apoptotic and cells under conditions of low confinement (5 μm). Cells that are under low confinement readily bleb, but do not normally migrate (Fig. 9B). For apoptosis, we used primary Human Dermal Fibroblasts (HDF), a fraction of which spontaneously undergo apoptosis upon serum withdrawal (Fig. 9A). Comparing apoptotic (HDF) and cells under low confinement (A375-M2; 5 μm), it was apparent that the blebs formed by apoptotic cells were highly dynamic (Fig. 9C; *right*). In contrast, the blebs formed by cells under low confinement could be long-lived (Fig. 9C; *right*). Bleb sizes for apoptotic and low confinement cells were similar (Fig. 9C; *left*). Comparing the largest bleb for each condition, several shape descriptors, including average roundness, circularity, solidity, and aspect ratio could differentiate between the two conditions (Figure 9D). Among these, the largest bleb aspect ratio was the most different, with low confinement cells having the largest aspect ratio (Fig. 9D). Thus, while the largest bleb size for each condition was similar, various shape descriptors indicate that blebs formed by cells under low confinement adopt a more elongated shape and can be long-lived (Fig. 9D).

**Figure 9.**
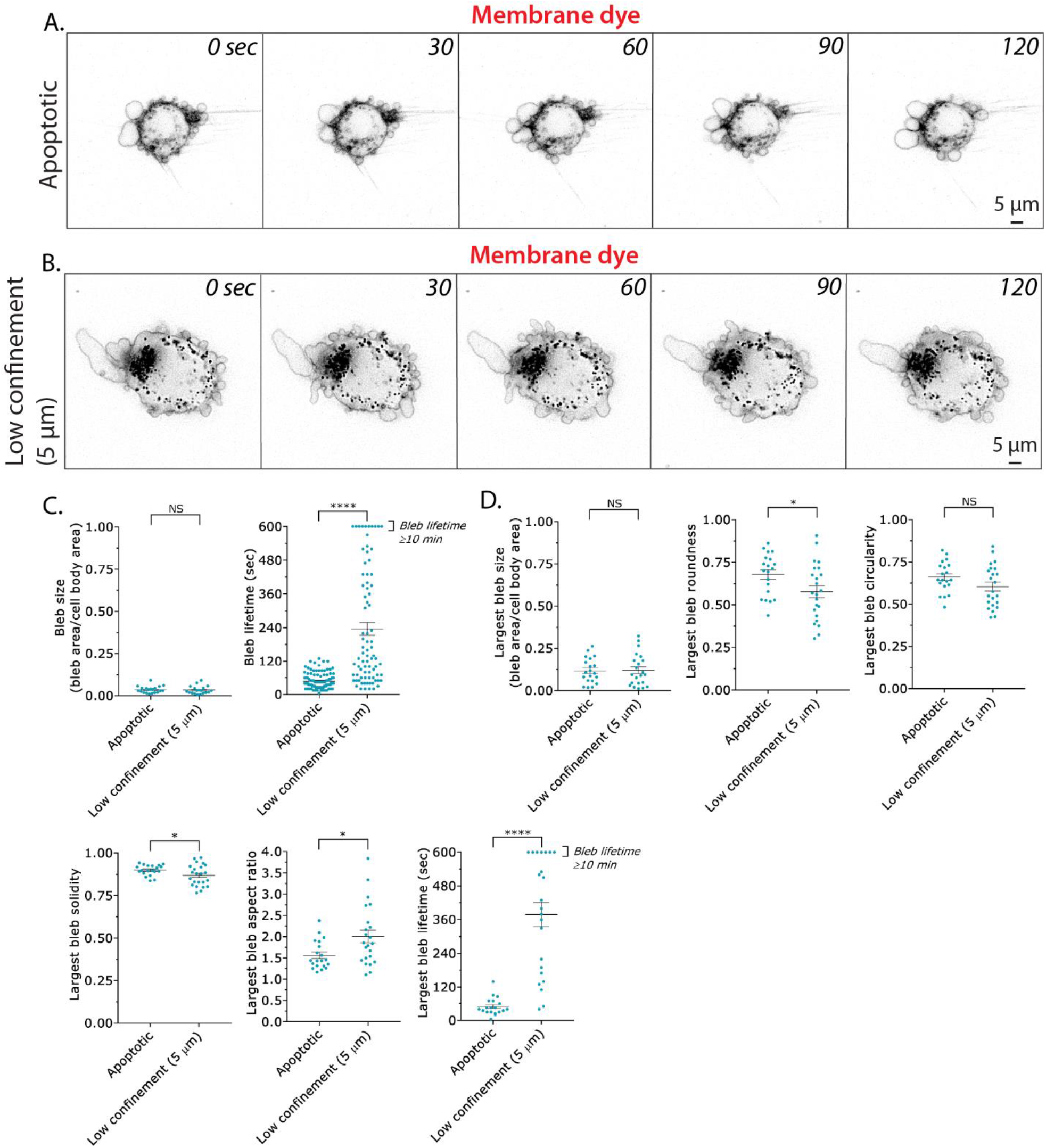
Analyze_Blebs may be used for measuring cell morphometrics under a variety of conditions. **A**. Montage of a primary Human Dermal Fibroblast (HDF). Prior to the start of the montage (∼20 min), cells were moved to serum free media to induce apoptosis. Cells were stained with membrane dye. **B**. Montage of an A375-M2 cell, which has been confined down to 5 μm. Under low confinement, cells are not normally motile. Cells were stained with membrane dye. **C**. Bleb size (*left*) and lifetime (*right*) for apoptotic (HDF) and low confinement (A375-M2; 5 μm) cells. Error is SEM. Statistical significance was determined using a two-tailed Student’s t-test. **D**. Largest bleb size (*left*), largest bleb roundness (*middle; top*), largest bleb circularity (*right; top*), largest bleb solidity (*left; bottom*), largest bleb aspect ratio (*middle; bottom*), and largest bleb lifetime (*right; bottom*) for apoptotic (HDF) and low confinement (A375-M2; 5 μm) cells. Error is SEM. Statistical significance was determined using a two-tailed Student’s t-test. All data are representative of at least three independent experiments. * - p ≤ 0.05, ** - p ≤ 0.01, *** - p ≤ 0.001, and **** - p ≤ 0.0001

Altogether, our results highlight the utility of Analyze_ Blebs in studying the morphometrics of diverse cell types under various conditions. However, these results should be interpreted with caution, as cell morphometrics may not be sufficient to fully understand the underlying basis for changes in the rate of LBBM.

## Discussion

The methods used to study or analyze cell migration continue to grow in sophistication. For instance, local surface curvature calculations with watershed segmentation, has been used to identify regions with high surface curvature as blebs [16]. Because of their unique shape, this approach is not likely to accurately identify leader blebs. Other versions of this software, which incorporates machine learning, may be able to automatically identify leader blebs [17]. A specific marker of leader blebs would also be helpful for increased automation. Fiji/ImageJ plugins, including ADAPT, also use measurements of curvature to track blebs [18, 19]. However, for accurate tracking, these plugins require imaging cells with very high temporal resolution in which the formation and disappearance of individual blebs is clearly resolved. While non-leader blebs often form and disappear in seconds, cell migration occurs over hours. Thus, analyzing fast amoeboid cells using these methods is problematic. These plugins are also able to measure changes in fluorescence, making it possible to determine when protein(s) are recruited to blebs [18, 19]. The Fiji/ImageJ-based plugin we describe here, Analyze_Blebs, was designed to require only minimal prior training in image processing. Using this plugin, researchers may rapidly obtain shape descriptors and cell migration parameters for the characterization of blebbing cells, including apoptotic and for cells undergoing fast amoeboid (leader bleb-based) migration. This also includes the analysis of nuclear blebs. Prior to Analyze_Blebs, all analyses, such as calculating the largest bleb area, were done entirely by hand – requiring a researcher to manually trace blebs within each frame of a time lapse. Here, using Analyze_Blebs, several shape descriptors and cell migration parameters may be determined simultaneously. Thus, Analyze_Blebs is anticipated to accelerate progress within this nascent field.

Using our previously described assay for the confinement of melanoma cells, we frequently observe three phenotypes, which we refer to as leader mobile (LM), leader non-mobile (LNM), and no leader (NL) cells. While the mechanistic basis for each phenotype is poorly understood, we report here that they may be characterized by certain shape descriptors. For instance, using Analyze_Blebs to rapidly assess several shape descriptors, we found that the average largest bleb area can differentiate between leader mobile (LM) and leader non-mobile (LNM) cells. Consequently, these data support our prior use of the largest bleb size as a meaningful shape descriptor [3, 10, 11]. Additionally, we report that average bleb size (i.e., all blebs), cell body size, whole cell roundness, and whole cell aspect ratio can differentiate between leader non-mobile (LNM) and no leader (NL) cells, as well as between leader mobile (LM) and no leader (NL) cells. Therefore, shape descriptors may be used to identify certain populations of cells under mechanical confinement.

Fast amoeboid migration depends on the rapid flow of cortical actin in leader blebs. Therefore, using live markers of F-actin that minimally perturb the natural behavior of cells is critical. Using Analyze_Blebs, we compared two commonly used live markers of F-actin, green and red fluorescent protein fusions of F-tractin and LifeAct [20, 21]. Qualitatively, we found each marker to similarly label the cortical actin in live cells. However, a comparison of migration parameters revealed that EGFP-F-tractin lowered the percentage of and speed of leader mobile (LM) cells. Similarly, transfection of FusionRed-F-tractin lowered the percentage and speed of leader mobile (LM) cells. Surprisingly, FusionRed alone had similar effects. We speculate that at least in part, this effect may be caused by an increase in free radicals, which are known to be generated by red fluorescent proteins in particular [22, 23]. To understand this phenomenon further, we used Analyze_Blebs to compare shape descriptors. Using this approach, we found that nearly all conditions led to an increase in the largest bleb size. Subsequently, we determined if the total level of F-actin in these cells was perturbed. For both green and red fluorescent protein fusions of F-tractin and LifeAct, the total level of F-actin was increased. Additionally, the transfection of FusionRed alone led to an increase in F-actin; therefore, FusionRed may have (undesirable) pleiotropic effects on cell physiology. Finally, through a cell compressibility assay, we measured cell stiffnesses. This revealed that transfection of EGFP-F-tractin, FusionRed alone, or FusionRed-F-tractin will increase cell stiffness. Importantly, an increase in cell stiffness perfectly correlated with inhibited migration. Therefore, shape descriptors must be interpreted with caution, as LBBM is also heavily regulated by cell mechanics [3, 10, 11].

Altogether, our data help to highlight some of the challenges a researcher faces when picking an F-actin marker. The most appropriate marker is likely to depend on the question being asked. For instance, because of its low affinity for G-actin, F-tractin may provide the highest contrast and so may be the best choice for high-resolution imaging [20, 24]. On the other hand, LifeAct may be the best choice if preserving specific cell behaviors is most important. Based on our data, mEmerald- or mScarlet-LifeAct are ideal if maintaining a high rate of leader mobile (LM) cells is critical. In conclusion, through the process of validating Analyze_Blebs, we have identified several important limitations of F-actin markers for the study of fast amoeboid (leader bleb-based) migration.

## Supporting information

Detailed protocol for using Analyze_Blebs

Troubleshooting table

Sample data

Movie 1 - Installation and Demo

Movie 2 - Additional Features and Troubleshooting

Movie 3 - Multi-cell and Individual Bleb Analysis

## Supporting Information

Supporting Information, which can be found online with this article, includes instructional videos, plugin code, sample data, detailed protocol for using Analyze_Blebs, and troubleshooting table. The most current version of the plugin can be found on GitHub (https://github.com/karlvosatka/analyze_blebs).

## Materials and Methods

### Cell culture and transfection

A375-M2 (CRL-3223) were obtained from the American Type Culture Collection (ATCC; Manassas, VA). Human Dermal Fibroblasts (HDF) were a gift from Dr. Paula J. McKeown-Longo (Albany Medical College). Cells were cultured in high-glucose DMEM (cat no. 10569010; Thermo Fisher, Carlsbad, CA) supplemented with 10% FBS (cat no. 12106C; Sigma Aldrich, St. Louis, MO), antibiotic-antimycotic (cat no. 15240112; Thermo Fisher), and 20 mM HEPES at pH 7.4 for up to 30 passages.

### Confinement

This protocol has been described in detail elsewhere [12]. Briefly, PDMS (cat no. 24236-10; Dow Corning 184 SYLGARD) was purchased from Krayden (Westminster, CO). 2 mL was cured overnight at 37 °C in each well of a 6-well glass bottom plate (cat no. P06-1.5H-N; Cellvis, Mountain View, CA). Using a biopsy punch (cat no. 504535; World Precision Instruments, Sarasota, FL), an 8 mm hole was cut and 3 mL of serum free media containing 1% BSA was added to each well and incubated overnight at 37 °C. After removing the serum free media containing 1% BSA, 300 μL of complete media containing trypsinized cells (250,000 to 1 million) and 2 μL of 3.11 μm beads (cat no. PS05002; Bangs Laboratories, Fishers, IN) were then pipetted into the round opening. The vacuum created by briefly lifting one side of the hole with a 1 mL pipette tip was used to move cells and beads underneath the PDMS. Finally, 3 mL of complete media was added to each well and cells were recovered for ∼60 min before imaging.

### Plasmids

FusionRed-F-tractin has been previously described [3]. EGFP-F-tractin (no. 58473), mEmerald (no. 53975), LifeAct-mEmerald (no. 54148), mScarlet-I (no. 85044), and LifeAct-mScarlet-I (no. 85054) were obtained from Addgene (Watertown, MA). 1 μg of plasmid was used to transfect 750,000 cells in each well of a 6-well plate using Lipofectamine 2000 (5 μL; Thermo Fisher Scientific) in OptiMEM (400 μL; Thermo Fisher Scientific). After 30 min at room temperature, plasmid in Lipofectamine 2000/OptiMEM was then incubated with cells in complete media (2 mL) overnight.

### Pharmacological treatments

Latrunculin-A (cat no. 3973) was purchased from Tocris Bioscience (Bristol, UK). DMSO (Sigma Aldrich) was used to make 5 mM stock solutions of Latrunculin-A. To disassemble actin, cells resuspended in flow buffer were treated with 5 μM Latrunculin A for 10 min at room temperature before flow cytometry.

### Flow cytometry

Roughly 1 × 10^6^ trypsinized cells in flow buffer (HBS with 1% BSA) were fixed using 4% paraformaldehyde (cat no. 15710; Electron Microscopy Sciences, Hatfield, PA) for 20 min at room temperature. After washing, cell pellets were resuspended in flow buffer and incubated with 0.1% Triton X-100, Alexa Fluor 647-conjugated phalloidin (cat no. A22287; Thermo Fisher), and DAPI (Sigma Aldrich) for 30 min at room temperature. Data were acquired on a FACSCalibur (BD Biosciences, Franklin Lakes, NJ) flow cytometer. Flow cytometric analysis was performed using FlowJo (Ashland, OR) software.

### Cell stiffness assay

The protocol for the gel sandwich assay was used with minor modifications [4]. 6-well glass bottom plates (Cellvis) and 18 mm coverslips were activated using 3-aminopropyltrimethoxysilane (Sigma Aldrich) for 5 min and then for 30 min with 0.5% glutaraldehyde (Electron Microscopy Sciences) in PBS. 1 kPa polyacrylamide gels were made using 2 μL of blue fluorescent beads (200 nm; Thermo Fisher), 18.8 μL of 40% acrylamide solution (cat no. 161-0140; Bio-Rad), and 12.5 μL of bis-acrylamide (cat no. 161-0142; Bio-Rad) in 250 μL of PBS. Finally, 2.5 μL of Ammonium Persulfate (APS; 10% in water) and 0.5 μL of Tetramethylethylenediamine (TEMED) was added before spreading 9 μL drops onto treated glass under coverslips. After polymerizing for 40 min, the coverslip was lifted in PBS, extensively rinsed and incubated overnight in PBS. Before each experiment, the gel attached to the coverslip was placed on a 20 mm diameter, 2 cm high PDMS column for applying a slight pressure to the coverslip with its own weight. Then, both gels were incubated for 30 min in media before plates were seeded. After the bottom gels in plates was placed on the microscope stage, the PDMS column with the top gel was placed on top of the cells seeded on the bottom gels, confining cells between the two gels. After 1 hr of adaptation, the height of cells was determined with beads by measuring the distance between gels, whereas the cell diameter was measured using a far-red plasma membrane dye (cat no. C10046; Thermo Fisher). Stiffness was defined as the height (*h*) divided by the diameter (*d*).

### Microscopy

Live high-resolution imaging was performed using a General Electric (Boston, MA) DeltaVision Elite imaging system mounted on an Olympus (Japan) IX71 stand with a computerized stage, Ultimate Focus, environment chamber (heat, CO2, and humidifier), ultrafast solid-state illumination with excitation/emission filter sets for DAPI, CFP, GFP, YFP, and Cy5, critical illumination, Olympus PlanApo N 60X/1.42 NA DIC (oil), UPlanSApo 20X/0.75 NA DIC (air), and UPLSAPO 10X/0.4 NA DIC (air) objectives, Photometrics (Tucson, AZ) CoolSNAP HQ2 camera, proprietary constrained iterative deconvolution, and vibration isolation table.

### Cell migration

To perform cell speed and plot of origin analyses, we used an Excel (Microsoft, Redmond, WA) plugin, DiPer, developed by Gorelik and colleagues and the Fiji plugin, MTrackJ, developed by Erik Meijering for manual tracking [25]. Brightfield imaging was used to confirm that beads were not obstructing the path of a cell. Cells that displayed directionally persistent migration over at least 5 frames (40 min) were classified as leader mobile (LM). Any cell with a large bleb that remained stable for at least 5 frames (40 min) was considered to have a leader bleb.

### Bleb morphology and dynamics

Roughly half of freshly confined cells were analyzed by hand and plugin. Cells analyzed by hand were traced from high-resolution images with the free-hand circle tool in Fiji (https://imagej.net/Fiji). Cells analyzed by plugin were processed according to the protocol attached to this paper, with data exported to Excel (Microsoft, Redmond, WA). Whole cell, cell body, and the largest bleb were traced on each frame. Measures including cell shape descriptors, percentage of cell area for leader blebs and percentage of cell area for total blebs were calculated in Excel (Microsoft). Frame-by-frame measurements were then used to generate an average for each cell.

### Statistics

All box plots are Tukey in which “+” and line denote the mean and median, respectively. Sample sizes were determined empirically and based on saturation. As noted in each figure legend, statistical significance was determined by either a two-tailed Student’s t-test or multiple-comparison test post-hoc. Normality was determined by a D’Agostino & Pearson test in GraphPad Prism. * - p ≤ 0.05, ** - p ≤ 0.01, *** - p ≤ 0.001, and **** - p ≤ 0.0001

### Data availability

The data that support the findings of this study are available from the corresponding author, J.S.L., upon reasonable request.

### Ethics Statement

This submission does not require an ethics statement.

## Acknowledgements

We would like to thank all members of the Logue Lab for their input throughout the development of the plugin. This work was supported by start-up funds from the Albany Medical College, a Young Investigator Award from the Melanoma Research Alliance (MRA; award no. 688232) (DOI: https://doi.org/10.48050/pc.gr.91570), and a Cancer Research Scholar Grant from the American Cancer Society (ACS; award no. RSG-20-019-01 - CCG) to J.S.L.

## Author Contributions

J.S.L. conceived and designed the study. S.B.L. performed all cell stiffness assays and contributed to plugin testing. K.W.V. performed all other laboratory research and was responsible for plugin development. J.S.L. wrote the manuscript with comments from K.W.V. and S.B.L.

