## Supporting Information for "A novel Fiji/ImageJ plugin for the rapid analysis of blebbing cells"

**Sample data.** Time-lapse imaging of a melanoma A375-M2 cell confined down to 3 μm. Membranes were labeled with a far red fluorescent dye. Time interval between frames is 8 min, 30 frames are provided in the form of a TIFF file. Each pixel is 0.1075 μm.

**Instructional video 1.** This tutorial video demonstrates how to install the Analyze_Blebs plugin along with the "ResultsToExcel" plugin, an essential add-on. It also demonstrates a typical run of the plugin on a blebbing cell.

**Instructional video 2.** This tutorial video highlights some of the more advanced image correction tools available prior to the analysis process. It also describes the different options for data output and how to troubleshoot using the plugin options menu.

**Instructional video 3.** This tutorial video demonstrates the more advanced features of the plugin, specifically multi-cell and individual bleb analysis.
