## Supplementary material for "A novel Fiji/ImageJ plugin for the rapid analysis of blebbing cells": Detailed protocol for using Analyze_Blebs

**Equipment**

- Fiji/ImageJ version 1.53d or later
- "ResultsToExcel" ImageJ Plugin
- Recommend use of DiPer Excel macros (Gorelik and Gautreau, 2014)

*"ResultsToExcel" Installation Guide*

1. In the Fiji/ImageJ Taskbar, go to "Help > Update…" to open the Plugin Updater
    a. Note: Be sure you are not selecting "Help > Update ImageJ…". That option deals with upgrading the base software of ImageJ, as opposed to the plugin updater.
2. When a window entitled "ImageJ Updater" appears, click the "Manage update sites" button.
3. Scroll down through the update sites and click on the checkbox next to "ResultsToExcel"
    a. If you do not see the ResultstoExcel option, first click on the "Update URLs" button
    b. If you still cannot find the proper site, click "Add update site", then enter "ResultsToExcel" in the "Name" box and the following URL into the URL box: https://sites.imagej.net/ResultsToExcel/

*"Analyze_Blebs" Installation Guide*

1. Ensure you have the latest version of the plugin downloaded from our GitHub page (https://github.com/karlvosatka/analyze_blebs).
2. In the Fiji Taskbar, go to "Plugins > Install…"
3. Navigate to the recently downloaded Analyze_Blebs.ijm file and select okay to install it.
4. Find a save location for the plugin. It is recommended that you install the plugin in the "plugins" folder for the FIJI software (…/Fiji/plugins).
5. Restart Fiji by closing it and re-opening it.
6. To run the plugin after installation, scroll to the bottom of the "Plugin" menu from the Fiji taskbar. You should find the file listed there.

**Standardizing image input and settings**

1. Open the time course data file of interest in Fiji/ImageJ as a hyperstack in Default color mode. Alternatively, open a plugin output file if desired to restart the plugin process from a later stage.
2. Enter the name that you would like prepended to each plugin output file in the box labelled "Data File Name:". The name of the chosen image file is filled in by default.
3. Indicate into which folder analysis files should be saved by clicking the "browse" button next to the box labelled "Data File Folder:". The file path of the chosen image file is filled in by default.
4. Indicate whether to export all data into an Excel document that corresponds to multiple data files from the same experiment. The data outputs corresponding to the chosen image will all be stored in a tab of this workbook. The most recently chosen Excel file for this

purpose will be filled in by default. Close the chosen Excel file before proceeding past this menu.

5. Indicate whether or not to employ the previously-chosen Z-isolation method and micron conversion settings.  If selected, the user will proceed through the Z-isolation method chosen previously (e.g., deciding on a single Z-slice vs. combining Z-slices).  Similarly, if chosen, the plugin will use the most recently chosen conversion rate for pixels to μm. If left unchecked, the plugin will offer all Z options and, if the image is not scaled to microns, will prompt the user to enter a conversion rate.
6. Indicate whether to analyze multiple cells on the same image.
7. Indicate whether to analyze individual blebs on this image. This is only recommended in cases where one can be confident that the full lifetime of the bleb can be observed from formation to dissolution.
8. Indicate whether to alter the plugin options.  The most recently used set of measurement options are used by default.  Upon first use of the plugin, the default measurements calculated for each output data file are area, centroid, perimeter, shape descriptors, Feret's diameter, and stack position.  Plugin options include changing measurement options or default sizes of cells or blebs, toggling an invert checker in case of errors with binary images, toggling automatic positioning of windows, and altering the default orientation of windows during tracing steps.
9. If the original image had multiple color channels, the colors will be split into different windows here.  Select the desired color channel in the drop down menu and click okay, and that channel will be used going forward.
10. If the image has multiple Z frames and step 5's box is unchecked, a prompt will appear to confirm the desired Z-stack isolation method: choosing a single Z slice or Z-projection.
11. To isolate a single Z slice, select the window with the best focus for the cell from the drop down menu and click okay on the dialog box.
12. To create a Z-projection, choose one of the options available from ImageJ for combining Z slices.  The default choice is a maximum intensity projection.
13. If the original image is scaled to pixels and the box in step 5 was not checked, enter the μm/pixel conversion rate specific to the microscope used to acquire the time series.
14. If multiple cells are to be analyzed on the image series, indicate the number of cells, the image to be used as a reference moving through the time series (e.g. tracer or a plugin output file like threshold) and whether to enter custom labels.
15. If custom labels for multiple cells are desired, enter the labels for each cell in the preferred analysis order.  This label will be appended to the data name chosen for the cell in each time series and the data for different cells will be saved in separate folders within the chosen save location.

**Edit the raw image series for reference**

16. Peruse the time series to determine if the frame range should be reduced to only those frames of interest, or if there are large objects with sizes comparable to the size of the cell that might interfere with thresholding and should be removed. If neither is true, move to step 21.
17. If the frame range needs to be edited, enter the first and last slice you would like to keep in the time frame and the interval of frames to keep (e.g., every 2nd frame).
18. If very bright objects with area comparable to the cell are present, remove the objects. If not, move to step 21. If multiple cells of similar brightness are present, move to step 21.
19. Hold the alt key and click on a spot that is representative of background on the first frame of the time series.
20. Circle all large, bright, and unwanted objects on each time frame and press the delete key to remove them.
21. If on any given frame the background color needs to be altered, indicate so in the "Remove Large Objects" window and repeat the step listed above for selecting the background color.
22. The plugin automatically creates and saves a TIFF image file called "Tracer.tif" that reflects your edits to the original data file and is used elsewhere in the plugin as a reference image for the original image series.
23. If alternate measurement options were chosen in step 6, you will be prompted to set new measurement parameters now from the set of options available in the "Set Measurements" dialog from ImageJ.

**Convert the image into a binary mask**

24. Review the preview of available thresholding methods for binarizing the image and choose your preference from the drop-down menu.
25. Check all frames for your chosen thresholding method, and then choose whether to proceed with this threshold method, try a different one, or manually draw the cell. You will be given the option to fill any holes that are present in the binary image in the next steps, so if there are fully contained holes present, proceed. You will also be given an opportunity to close any holes or remove any noise attached to the image, so if the noise is not excessive and there are not many large holes to close, proceed.
26. If after reviewing the above a different method is desired, return to step 24. If a proper threshold is established, move to step 30.
27. If unable to find a threshold method that produces even a fairly clean image of the cell, choose the "Draw the cell" option.
28. To begin drawing the cell, click the "Synchronize All" button in the "Synchronize Windows" window as directed.
29. Trace the outline of the cell on the tracer window, make sure "Draw the cell" option is selected, then press okay.

30. Repeat for every frame of the image as needed, and when finished, choose “Time course is complete” and press okay. Move to step 46.

**Make corrections to the mask**

31. Check the time course to see if any white holes are present inside the black shape of the cell . If holes are noted, they will be filled. If not, move to step 36.
32. If holes were filled initially, check the image again for any remaining holes or edges of the cell that are not closed and indicate appropriately. If not, move to step 36.
33. Begin the process of filling remaining holes by clicking the “Synchronize All” button in the “Synchronize Windows” window.
34. On each frame, trace a line that starts and ends in the black, filled in area of the cell so that the space missing from the cell is enclosed in an outline. This creates a new hole that is still part of the cell.
35. Press okay with the “Draw the cell outline” button selected, and the new cell outline will be added and filled. If the outline you drew didn’t fill in properly, then you didn’t make a closed shape and you need to ensure the outline is closed within the cell.
36. Complete hole filling for the time course, and when finished, indicate so and proceed. In the case of multiple cells, if preferred, ensure all interesting cells have no holes at this step.
37. Choose the minimum size cut off for the smallest object in frame that will be counted as a cell, in $\mu m^2$. The default is 200 $\mu m^2$, chosen based on observations from A375-M2 cells. All objects still in frame, including any background noise, that are below this size will be removed.
38. Check the time course to see if any noise is attached to the cell(s) of interest, including two cells touching each other, and indicate so. If not, move to step 44.
39. Begin the process of manually separating noise by clicking the “Synchronize All” in the “Synchronize Windows” window.
40. On each frame, trace a line that starts and ends in the white background and that properly outlines the cell at the point where the noise is attached.
41. Press okay with the “Separate noise object” button selected, and the object will be separated from the cell. If the shape is still there, it is either larger than your cell size in step 36 or is still attached to the cell of interest.
42. Complete for the time course, and when finished, indicate so and proceed.
43. If multiple cells were identified in step 13, identify which of them are present on the current threshold image based on their labels.
44. In the case of multiple cells, any cells not present on the current threshold will need to either be re-thresholded or re-drawn. If the cell described in the dialog is present, you will be able to proceed without threshold changes. There is also an option to change the frame range for a particular cell following the process outlined in step 16.
45. Click “Unsynchronize All” in the “Synchronize Windows” window.

46. For every frame where you are prompted to do so, click on the blue number corresponding to the cell of interest, and ensure that cell has a light blue outline. Try not to drag on the number at all, as small errors in measurement will occur if the blue outline does not accurately cover the edge of the red cell shape. Click okay in the dialog box, and all other cells will be removed from that frame.
47. The plugin automatically measures the whole cell binary image and saves the image (.tif) and data (.csv) as the "whole_cell" measurement series.

**Identify the cell body to isolate blebs**

48. Click the "Synchronize All" button in the "Synchronize Windows" window.
49. Circle the cell body so that blebs lie outside of the circle. There is no need to follow the cell outline, except along the path where the cell body and blebs meet.
50. When done with a frame, ensure the "Remove the cell body" option is checked, then press okay. The cell body will be removed from the cell_mask window. You can continue to edit the same frame or move to others as desired.
51. Repeat steps 47-49 for the time course and when finished, indicate so and press okay.
52. Choose the minimum size cut off for the smallest object in frame that will be counted as a bleb, in $\mu m^2$. The default is 10 $\mu m^2$, chosen based on observations from A375-M2 cells. All objects still in frame, including any background noise, that are below this size will be removed.
53. The plugin automatically measures the mask and saves the image (.tif) and data (.csv) as the "all_blebs" measurement series.
54. The plugin automatically creates a series for the cell body and the largest bleb in each frame, creating files as above. (Landini *et al.*, 2008)

**Track Individual Blebs**

55. If the individual bleb analysis box was selected in step 7, the individual bleb analysis menu will appear.
56. Select the number of the bleb you would like to analyze. The plugin automatically counts sequentially from 1. If you are running analysis of more blebs after a previous run, make sure you change the number of the bleb to a number greater than your last analyzed bleb to ensure data is not overwritten.
57. Compare the tracer and all_blebs images to find the frame range for the bleb of interest. If Synchronize Windows is open, you can use it to make this easier, but you will have to un-synchronize later.
58. One satisfied, click okay to proceed, and click un-synchronize windows in the Synchronize Windows box as prompted.
59. Select the bleb of interest on each frame when prompted to do so by clicking on the blue number corresponding to the bleb. Try not to drag this outline and instead ensure it

matches the outline of the red bleb shape. Once the outline turns light blue and is in place, click okay.

60. Repeat step 59 throughout the time series and return to step 55, repeating as necessary. Once finished, click the check box to indicate so and proceed.

**Multiple Cells**

61. If multiple cells are to be analyzed, return to step 43 and repeat as needed.

Landini G. Advanced shape analysis with ImageJ. Proceedings of the Second ImageJ User and Developer Conference, Luxembourg, 6-7 November, 2008. p116-121. ISBN 2-919941-06-2. Plugins available from https://blog.bham.ac.uk/intellimic/g-landini-software/

Gorelik, R., and Gautreau, A. (2014). Quantitative and unbiased analysis of directional persistence in cell migration. Nat Protoc *9*, 1931–1943.
