## Supplementary material for "A novel Fiji/ImageJ plugin for the rapid analysis of blebbing cells": Troubleshooting table

| Step | Problem | Possible reason | Solution |
|---|---|---|---|
| 1 | Trying to rerun the plugin from previous output files doesn't seem to work | Using the wrong plugin output files | The plugin will not operate properly from cell body or largest bleb images. To recalculate from a copy of the raw data with bright objects removed and the preferred frame range selected, start the plugin with the "tracer" image and proceed to choosing a threshold. Start from "_threshold" to edit it by removing noise and filling holes in the cell. To again separate the cell body and blebs, start from "_whole_ cell". To re-calculate "_cell_body" and "_largest_bleb" images, open "_whole_cell and "_all_blebs" images to start. All images past the tracer step require the tracer image to be open; the user may be prompted to open it if it isn't open already. |
| 4 | Some or all of the cell info isn't saved properly into the Excel file with multiple cells saved from one experiment (or IF.error.FileNotFound error appears) | The experimental Excel file is still open | Close the experimental Excel file and the rest of the information from the plugin running should load into the file properly. You can add any missing data from earlier in the process back in using the .CSV files output in your data folder. |
| 4 | In the optional Excel experimental summary file, multiple seemingly redundant entries appear in the sheet for one cell (e.g. there is a whole_cell category listed twice, perhaps with different data each time) | The plugin was run repeatedly using the same data name and saved data from multiple runs in the same sheet | Be sure to change the name used for each set of data if you want it to appear in separate tabs. The data will be entered for each tab in the order in which the cell was processed, so later plugin runs with the same plugin name will appear farther on the right in the same tab. Refer to the "_excel" file and CSV files for each output image series to correct errors as needed, or simply delete the sheet for the cell from the Excel summary file and re-run the cell through the plugin. |
| 13 | Still received a prompt to change pixel conversion rate despite clicking on "Use previous settings" button in step 6 | Running the plugin for the first time with any file requiring pixel conversion | Enter the conversion rate and move forward. In future, the image should be converted with that conversion rate as needed. |
| 20, 32 | Clicking on the "remove/draw" button on the dialog box and then clicking forward in the time series is tedious | Auto-positioning feature is turned off. Or, screen is too small to see all windows well, and windows must be shuffled to use the plugin | First, check if auto-positioning is on in the plugin options menu. If you prefer to keep it off, you have other options. Consider moving the frame forward using the arrow key. If the image is selected, click the edge of the image window to select it, hold shift and click the arrow key to move forward or backward while still keeping the same area selected. This allows the user to delete a |

| | | | |
|---|---|---|---|
| | | | non-moving object from the frame quickly without redrawing the outline of that object. |
| 20 | It takes too long to remove large background objects from the raw image | User is attempting to remove objects that will be excluded by size later or that are not bright | Remember only to remove any very bright objects that are close to the size of the cell. Also, try removing objects by pressing delete instead of via the dialog box method. Do not remove other cells through this method unless they are much brighter than the cell of interest. Refer to the previous troubleshoot tip for keyboard controls that should speed up this step as well. |
| 25 | Attempted to remove large objects on the raw image in step 19, but on some frames, there are still large images showing up where the erase marks were made | Background painting color was not adjusted by user and photobleaching caused background color to darken | Re-run the original image file in the plugin, and while editing the raw image in steps 14-17, reset the background color every 5 frames or less using the “Reset Background” command in the dialog. |
| 25, 30 | The “_threshold” image has cells show up in white on a black background as opposed to the plugin-standard black cell on white background | Improper dimensional assignment of frames, or the cell is very large and covers several edges of the screen. | Turn on the inversion check feature, which can be accessed in the plugin options menu under the label “Invert Check”. Try toggling this setting as needed to prevent recurring instances of the problem. Remember to check the appropriate box in step 6 if you want to alter the inversion check feature. |
| 37, 52 | The cell or blebs disappears on some or all frames after proceeding through these size exclusion steps | The default size chosen for the cell or for blebs is incorrect | Enter a different size as desired at these two steps. After running repeated analysis and developing a sense of the typical size of the cell and of blebs in the condition studied, change the defaults through the plugin options toggled in step 6. |
| 46, 59 | Cursor rapidly switches between a plus shape and normal cursor shape when trying to click on blue numbers for cells/blebs | Windows are still synchronized | Click “Unsynchronize Windows” in the Synchronize Windows box. This should make it easier to select the outline for your bleb/cell of choice. |
| 49 | While tracing, two blebs adjacent to each other share a border and are not seen as separate in the whole cell mask image | The blebs are too close and bright to be identified as separate | Draw the outline of the cell body around the bottom of the blebs, then, at the point where the blebs should be divided, draw a perpendicular line towards the outside of the cell and trace the separating line between the blebs. Be sure this line reaches the outside black area, then trace the same line back in towards the cell body and continue tracing. The blebs will be separated by that line. |

| | | | |
|---|---|---|---|
| 53 | Many, many more blebs than expected appear in the data outputs for the "all_blebs" image | The bleb analysis size chosen in this step was zero or close to zero. | Choose a larger bleb cutoff size. Otherwise, noise objects that are present but are too small to see will be counted as blebs. Picking a size of zero fails to exclude pixel-sized noise objects. |
| 56 | Data from individual bleb analysis doesn't match expectations | Data was overwritten by not changing the number label for each bleb | Be sure to change the number for the bleb of interest if bleb analysis was run on a data file previously, or else old bleb files will be overwritten. |
| n/a | Large "Console" ImageJ error window with red text inside of it appears during running of plugin | None | These errors should not affect the running of the plugin. They occur most often when analyzing multiple cells on the same image series. You may ignore the window or close it and proceed. |
