## Supplementary figures and images for "A novel Fiji/ImageJ plugin for the rapid analysis of blebbing cells"

### Sample data

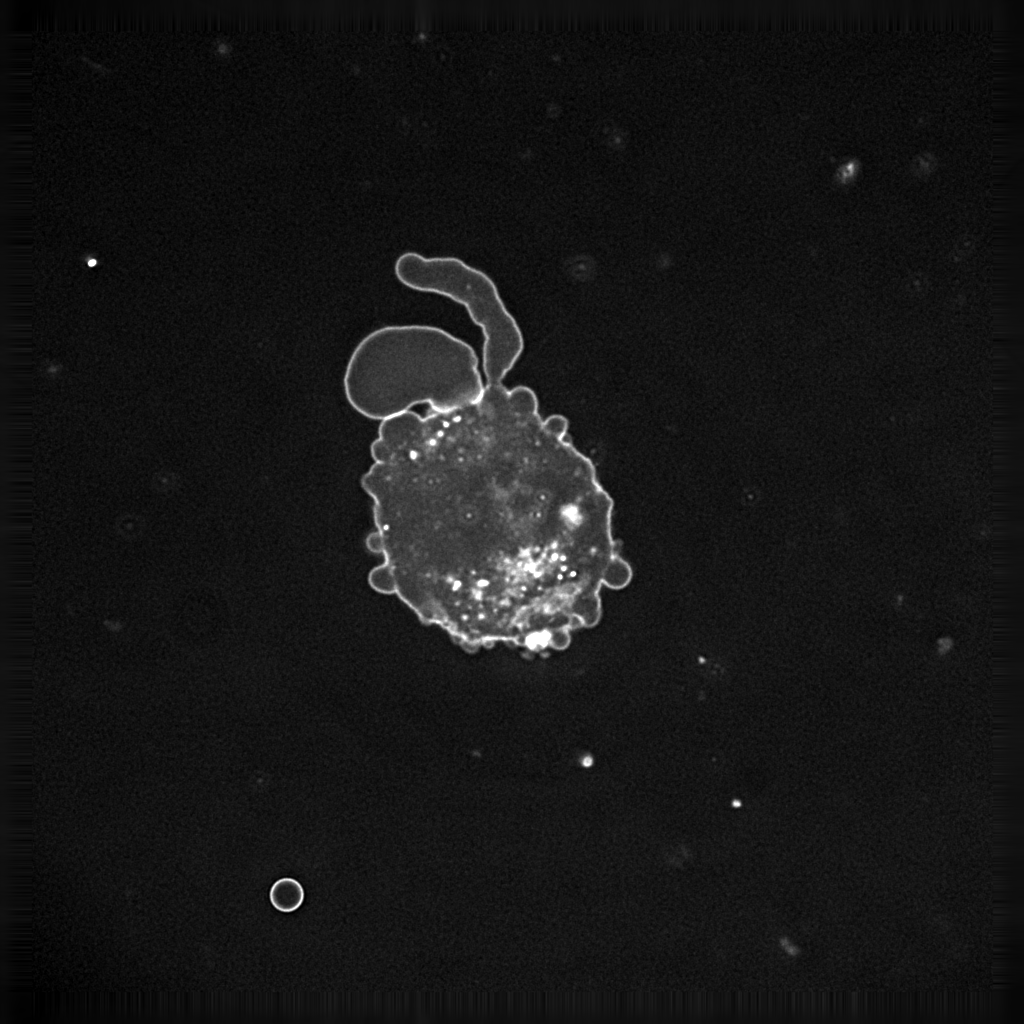
